# Synchrony and complexity in state-related EEG networks: an application of spectral graph theory

**DOI:** 10.1101/729806

**Authors:** Amir Hossein Ghaderi, Bianca R. Baltaretu, Masood Nemati Andevari, Vishal Bharmauria, Fuat Balci

## Abstract

To characterize differences between different state-related brain networks, statistical graph theory approaches have been employed to identify informative, topological properties. However, dynamical properties have been studied little in this regard. Our goal here was to introduce spectral graph theory as a reliable approach to determine dynamic properties of functional brain networks and to find how topological versus dynamical features differentiate between such networks. To this goal, 45 participants performed no task with eyes open (EO) or closed (EC) while electroencephalography data were recorded. These data were used to create weighted adjacency matrices for each condition (rest state EO and rest state EC). Then, using the spectral graph theory approach and Shannon entropy, we identified dynamical properties for weighted graphs, and we compared these features with topological aspects of graphs. The results showed that spectral graph theory can distinguish different state-dependent neural networks with different synchronies. On the other hand, correlation analysis indicated that although dynamical and topological properties of random networks are completely independent, these network features can be related in the case of brain generated graphs. In conclusion, the spectral graph theory approach can be used to make inferences about various state-related brain networks, for healthy and clinical populations.

**Author Summery:** By considering functional communications across different brain regions, a complex network is achieved that is known as functional brain network. Topologically, this network is constructed by different nodes (activity of brain regions or signals over recording electrodes) and different edges (similarity, correlation or phase difference between nodes). Paths, clusters, hubs, and centrality of nodes are examples of topological properties of these networks. However, synchrony and stability of functional brain networks can not be revealed by consideration of topological properties. Alternatively, spectral graph theory (SGT) can demonstrate the dynamic, synchrony and stability of graphs. But this approach has been studied little in brain network analysis. Here, we employed SGT, as well as topological methods, to investigate which approaches are more reliable to find differences between distinct state-related brain networks. On the other hand, we investigated correlations between topology and dynamic in different type of networks (brain generated and random networks). We found that SGT measures can clearly distinguish between distinct state-related brain networks and it can reveal synchrony and complexity of these networks. Also, results show that although dynamic and topology of random-generated graph are completely independent, these properties exhibit several correlations in the case of functional brain networks.

## 1 Introduction

After the development of graph theory analytical (GTA) tools in physics ^1–3^, these tools have been used to investigate the statistical aspects of functional brain networks ^4–6^. By exploiting this method, earlier studies focused on topological properties of functional brain networks to revealed the most primary features of the brain connectivity^7–11^, for example, in describing neural basis of brain functions/dysfunctions in health/disease ^6,12–14^. However, in the last decade, GTA (using more sophisticated measures) has also been used to unravel neural networks that are involved in different states of brain/mind and then associate the topological properties of these neural networks to further evaluate the brain connectivity at global or local levels ^7–11,15,16^.

Recently, studies using GTA to evaluate brain networks have been merged under the name of “*network neuroscience”*^8^. As the second important advance in the field, recent studies have been focused on dynamics of functional brain network ^17–26^. To this aim, most of these investigations employed a *sliding window* method (i. e. by dividing the entire signal into separate or overlapping parts) to investigate the dynamic functional connectivity ^17,22–26^. In the related literature, network connectivity that is derived from the entire signals (or averaging between sections) is referred to as ‘static functional connectivity’^26^, whereas, networks that are disclosed by the *sliding window method* are labeled under the term ‘dynamic functional connectivity’^26^. Thus, by employing either of these methods, the topological (static/dynamic) of functional networks can be revealed^22–26^.

In contrast, from a mathematical perspective, the topology of a network (association between nodes and edges^17^) is considered as a statistical (not dynamical) issue ^27–30^. Then the topological parameters, namely: shortest path, clustering coefficient and degree^8^ cannot be attributed to the dynamic properties of complex networks, such as, synchronization^27–29^ and level of coupling in a graph^31^. Therefore, even small-world graphs can be non-synchronized^31^ and two networks with same topology can exhibit two different levels of synchronization ^27^. From this point of view, the dynamical aspects of graph are invariant in relation to topology and a different mathematical method is required to describe that. To this aim, a specific filed in graph theory that is known as *“spectral graph theory, SGT”*^32^ was introduced. Mathematically, this approach adopted linear algebra to decompose adjacency or Laplacian matrix to the corresponding eigenvalues and eigenvectors^32^.

Despite the important role of the SGT to find the dynamic of functional brain networks, only few studies have implemented SGT to elucidate the dynamical properties of the brain^33–37^ in network neuroscience. These studies indicated that the overall coupling strength between functional brain network-units (brain regions or electrophysiological electrodes) can be changed by age^38^ or by disorders like schizophrenia^36^ and epilepsy^35,37^. However, the differences/correlations between results derived from SGT approach and topological methods are still unclear. On the other hand, most of previous studies have been performed on binary on sparse adjacency matrices^34,36,39,40^ or did not consider the methodological problems such as weight-conserving ^41^ of SGT measures that they applied on weighted graph ^37,38^.

Here, to find the associations between the results of spectral and topological method, and then clarify the efficiency of these approaches, we investigated a well-studied paradigm — what is the difference in electroencephalography (EEG) between eye-closed and eye-open conditions^42–44^? We analyzed state-related EEG network during resting state eyes-closed (rsEC) and resting state eyes-open (rsEO) conditions. Previous studies demonstrated that functional brain networks are changed during rsEC and rsEO conditions^42–45^. Thus, it is an appropriate example to compare the eligibility of both dynamical and topological approaches in network neuroscience. We also used a K-mean clustering approach with different inputs that are derived by topological/spectral methods. This approach can clarify whether spectral or topological graph measures can separate state-related EEG networks during two different resting state conditions. Furthermore, to figure out the ‘weight distribution conserving’ of topological and spectral approaches, we applied above mentioned methods to randomly generated matrices with same means and standard deviations but with different distribution of weights (exponential vs. normal distribution). Thus, we could find which of above approaches can demonstrate the randomness of networks even with different distribution of weights. Since the SGT reveals the dynamic rather than the topology of graph, we hypothesized that despite the topological approach, this method is invariant in relation to distribution of weights. By this way, independent information can be obtained by these two different approaches (topological and spectral).

Since the brain synchrony and complexity are frequently reported to be related to each other^20,46,47^, we considered the complexity of networks through the novel application of Shannon entropy, and, then the possible relation between the functional brain network complexity and synchrony was evaluated. We further hypothesized that there is an inverse relation between network complexity and synchrony while complexity is irrelevant to topology of network. To our knowledge, the relationship between SGT measures, Shannon entropy and topological graph indices has been investigated for the first time in this study. We found that although new information about functional brain networks can be revealed by SGT, still there are several correlations between SGT and topological measures in brain networks analysis.

## 2 Results

### 2-1. Probability distribution of weights

The probability distributions of four types of matrices (i.e., rsEO, rsEC, normal distribution, exponential distribution) are shown in Fig. 1. The results of the probability distribution analysis indicate that the brain-generated adjacency matrices, which are representative of brain coherence between all the electrodes, show a semi-exponential distribution. In addition, a power law constraint is established on the brain-generated connectivity weights.

**Figure 1:**
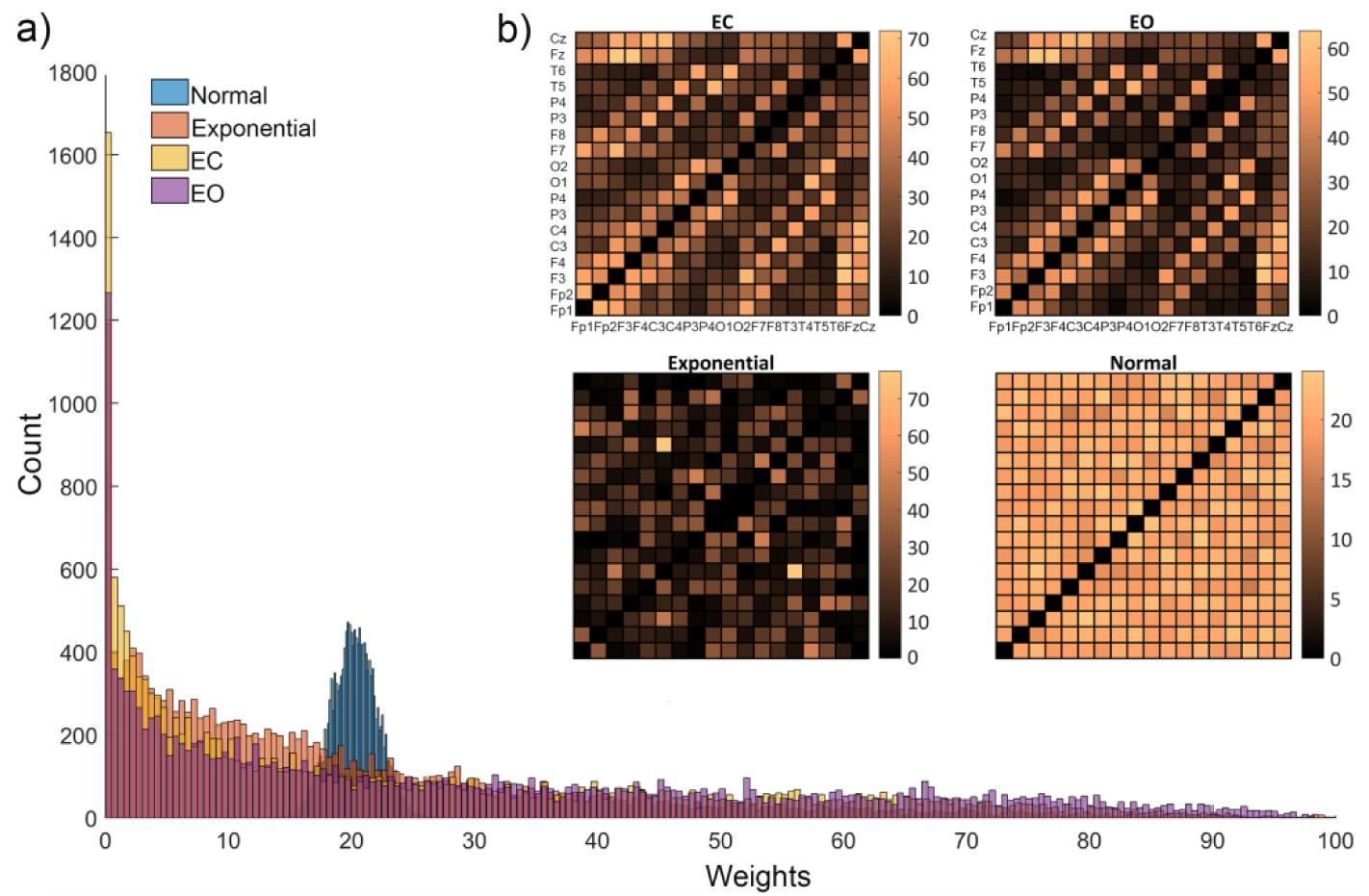
**a)** The probabitity distribution of weights in adjacency matrices. We generated two sets of random networks (45 networks with random exponential distribution and 45 networks with random normal distribution of wheights) using mean and SD of brain generated networks (rsEC and rsEO conditions). The rsEC and rsEO networks show semi-exponantial distribution while the normal distribution is completely different from the brain-generated distributions. **b)** Average of adjacency matrices (over 45 matrices for each subject). In the rsEC and rsEO matrices, each row/column represents an EEG channel (not applicable for random distribution matrices). Lighter colors are associated with higher coherences between channels, while, darker colors corrspond to lower coherences.

The R-squared value was also calculated between histograms. Results from this analysis indicate that considerable R-squared values were obtained for: 1) rsEO and rsEC conditions (0.883), 2) rsEC and random exponential (0.877), and 3) rsEO and random exponential (0.899). In contrast, the R-squared values between normal and exponential distributions (0.398), normal and EC distributions (0.474), and normal and EO distributions (0.412) exhibited a weak correlation.

### 2-2. Graph theoretical analysis

#### 2-2-1 Topological measures of weighted graph

False discovery rate (FDR) analysis results showed that there were significant differences between the two eye conditions in clustering coefficient (F=6.647, p-value < 0.0001) and characteristic shortest path (F = 3.749, p-value < 0.0001). Higher values of clustering coefficient and characteristic shortest path were found in our rsEC condition (Figs. 3, 4). Investigation of individual participant activity indicates that 40/45 participants exhibited the same pattern for the clustering coefficient, whereas 37/45 participants showed a similar pattern for characteristic shortest path (EC > EO). FDR analysis indicated that clustering coefficient and characteristic shortest path are significantly different for two random sets of graphs. The significant difference of clustering coefficient was observed between random graphs with exponential distribution and two brain-generated datasets (rsEC and rsEO). However, clustering coefficient of random graph set with normal distribution was not significantly different from rsEC condition. In characteristic shortest path, although significant differences were observed between random graphs with exponential distribution and two brain-generated graph sets (rsEC and rsEO), random graph with normal distribution exhibited significant difference only with the rsEC condition (Fig 2).

**Figure 2:**
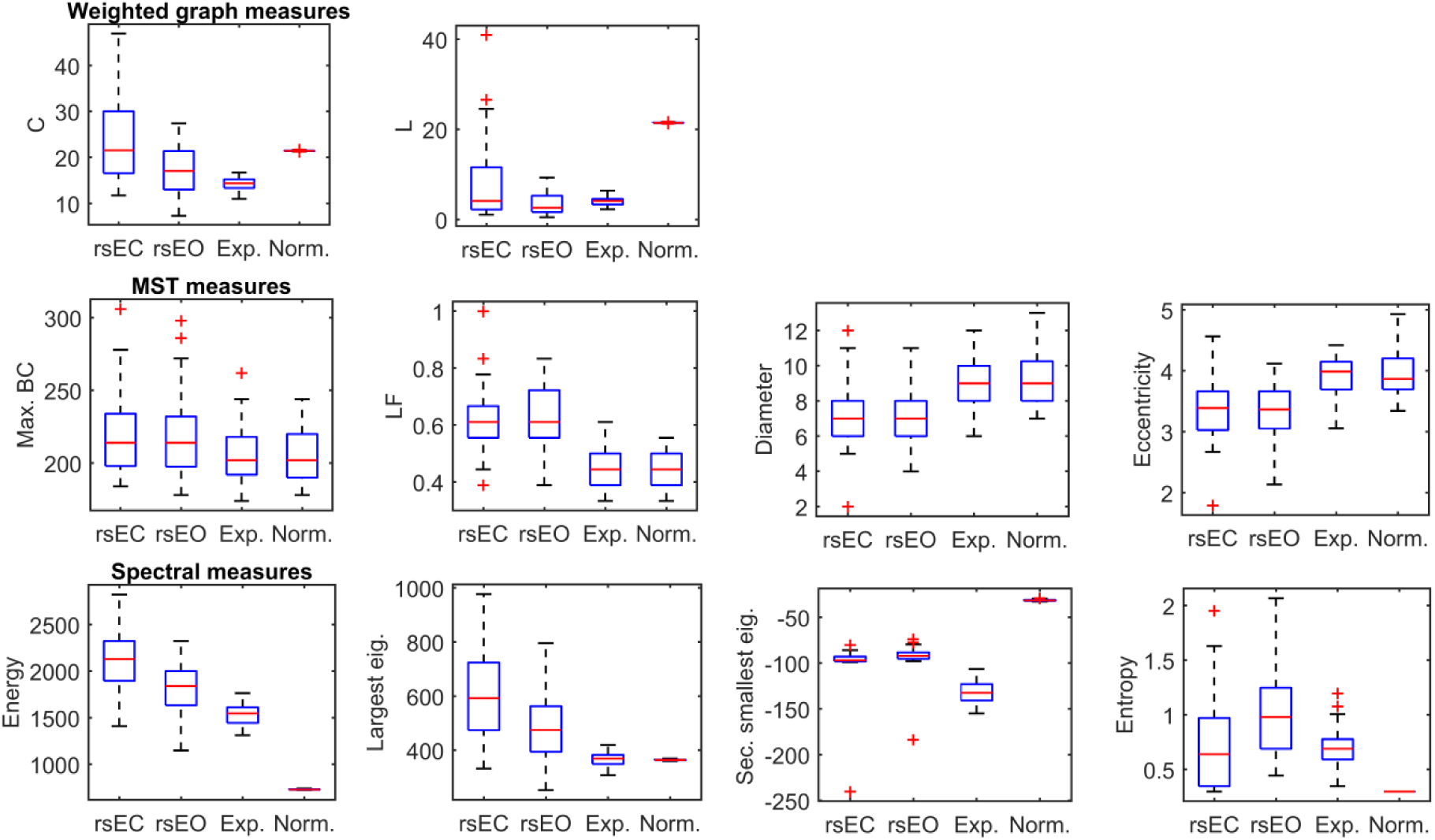
Weighted graph measures (top). The rsEC network exhibits higher value of C while the random exponential network shows the lowest value. C is sensitive to distribution but not to randomness. Random normal network shows the highest L, while, the rsEO and random exponential networks have lower values of L. L is sensitive to distribution of weights. MST measures (middle). Max BC, LF, diameter and eccentricity are sensitive to randomness but not to distribution. These measures cannot distinguish the rsEC and rsEO networks. Dynamical measures (below). The highest values of energy and largest eigenvalue are related to the rsEC network, whereas, the rsEO network shows the highest value of entropy. Energy is sensitive to distribution. The largest eigenvalue is sensitive to randomness but not to distribution. The second smallest eigenvalue exhibits same values for rsEC and rsEO networks but it can distinguish brain-generated and random networks.

#### 2-2-2 MST measures

After applying FDR correction to the results of a within-subject permutation t-test, we found no significant differences in betweenness centrality (F=0.0567, p-value=0.521), leaf fraction (F=-0.2633, p-value=0.354), eccentricity (F=0.333, p-value=0.355) and diameter (F=0.4997, p-value=0.308) between the rsEO and rsEC conditions. Individual results indicated that participants did not share the same pattern of results for these measures (Fig. 2). Further, we found that both random datasets showed significant differences from brain generated graphs (in all the MST measures). However, there were no significant differences between two randomly generated graphs.

#### 2-2-3 Dynamical Measures (Synchrony and Complexity)

After FDR analysis, significant differences between the two eye conditions (rsEO and rsEC) across the tested dynamical indices were observed. Significantly higher values of Shannon entropy were observed in the rsEO condition (F=-6.828, p-value=0.0001) while higher values of largest eigenvalue (F=-7.045, p-value=0.0001) and energy (F=- 10.852, p-value=0.0001) were observed in the rcEC condition (Fig. 2). However, there was no significant difference between rsEC and rsEO in the second smallest eigenvalue. The differences between random graphs were significant in energy, entropy and second smallest eigenvalue, but two random graph sets exhibited statistically no significant deference in largest eigenvalues. Significant differences were observed between brain-generated and random-generated graphs in the largest eigenvalue, the second smallest eigenvalue, and the energy. However, there was no significant difference between brain-generated and random-generated graph in entropy (Fig. 2).

#### 2-2-4 Correlation Analysis

Figure 3 indicates that several significant positive and negative correlations exist between the graph measures that we tested. According to the correlation analysis, in both of random networks, generally three main clusters were observed that exhibited high correlation. First, a cluster between the dynamical indices (largest eigenvalue, second smallest eigenvalue and energy). Second, a cluster between weighted graphs measures (L and C), and third, a cluster of MST measures (BC, LF, diameter and eccentricity). For these analyses, we did not find any significant correlation across these three clusters.

**Figure 3:**
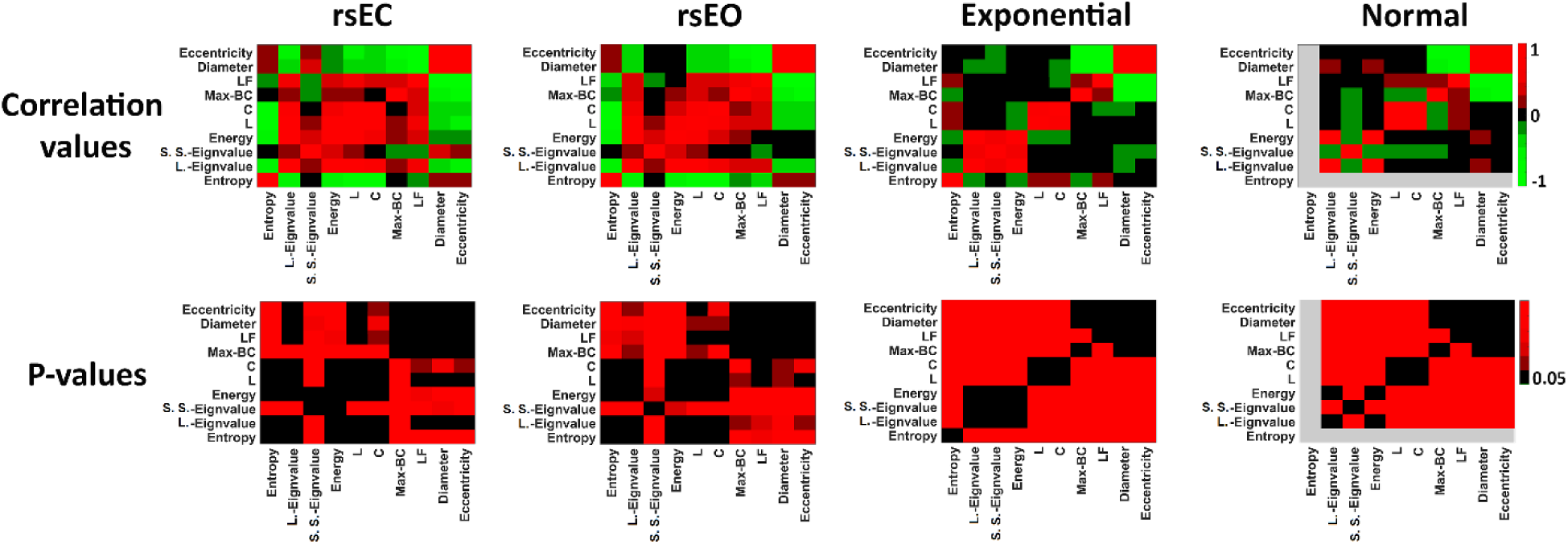
Heatmap of correlations between measures (Top). Light green colors are associated with negative high correlation between measures, while, light red color indicates positive high correlation between measures. Dark green, dark red and black indicate low correlation between measures. P-value of correlations after Bonferroni correction (Bottom). Black cells indicate significant correlation between measures (p<0.05) while red color indicates non-significant correlations (p>0.05). Different groups of correlation are observed for random networks with exponential and normal distribution of weights. In the correlation Heatmap of these networks, we can distinguish at least three groups for three types of measures; spectral, topological for weighted graph and MST. These groups are completely separated for random networks with no correlation across them. However, several correlations are observed across these groups for EEG networks (see discussion section). S. S eigenvalue is second smallest eigenvalue. L. eigenvalue is largest eigenvalue.

However, in the brain generated graphs (rsEC and rsEO) we found several significant correlations across dynamical, weighted and MST measures. In both conditions, significant positive correlations were observed between the maximum spectral measures (largest eigenvalue and energy) and the weighed graph measures (C and L). On the other hand, for both conditions, entropy exhibited a significant negative correlation with spectral measures (largest eigenvalue and energy) and weighed graph measures (C and L). In rsEC, there was a significant correlation between the largest eigenvalue and the MST measures (positive correlation with LF and negative correlation with diameter and eccentricity). Significant correlations between the weighted graph measures and the MST measures were also observed in both conditions. There was no significant correlation between the second smallest eigenvalue and other measures. All correlational measures as well as p-values (after Bonferroni correction) are presented in Fig. 3.

#### 2-2-5 Clustering Analysis

The scatter plots indicate that two-dimensional patterns (using both dynamical and topological measures) are not linearly separable (Fig. 4). Based on the results of the FDR analysis on the data from the two conditions, we used the measures that had significant differences between them, as inputs for the linear k-means approach. The input patterns differed across the dimensions, which ranged from two to five. All possible combinations of inputs were tested and incorrect assignments for each model were determined. The best accuracy (93.3%) was obtained using clustering coefficient, characteristic shortest path, and entropy.

**Figure 4:**
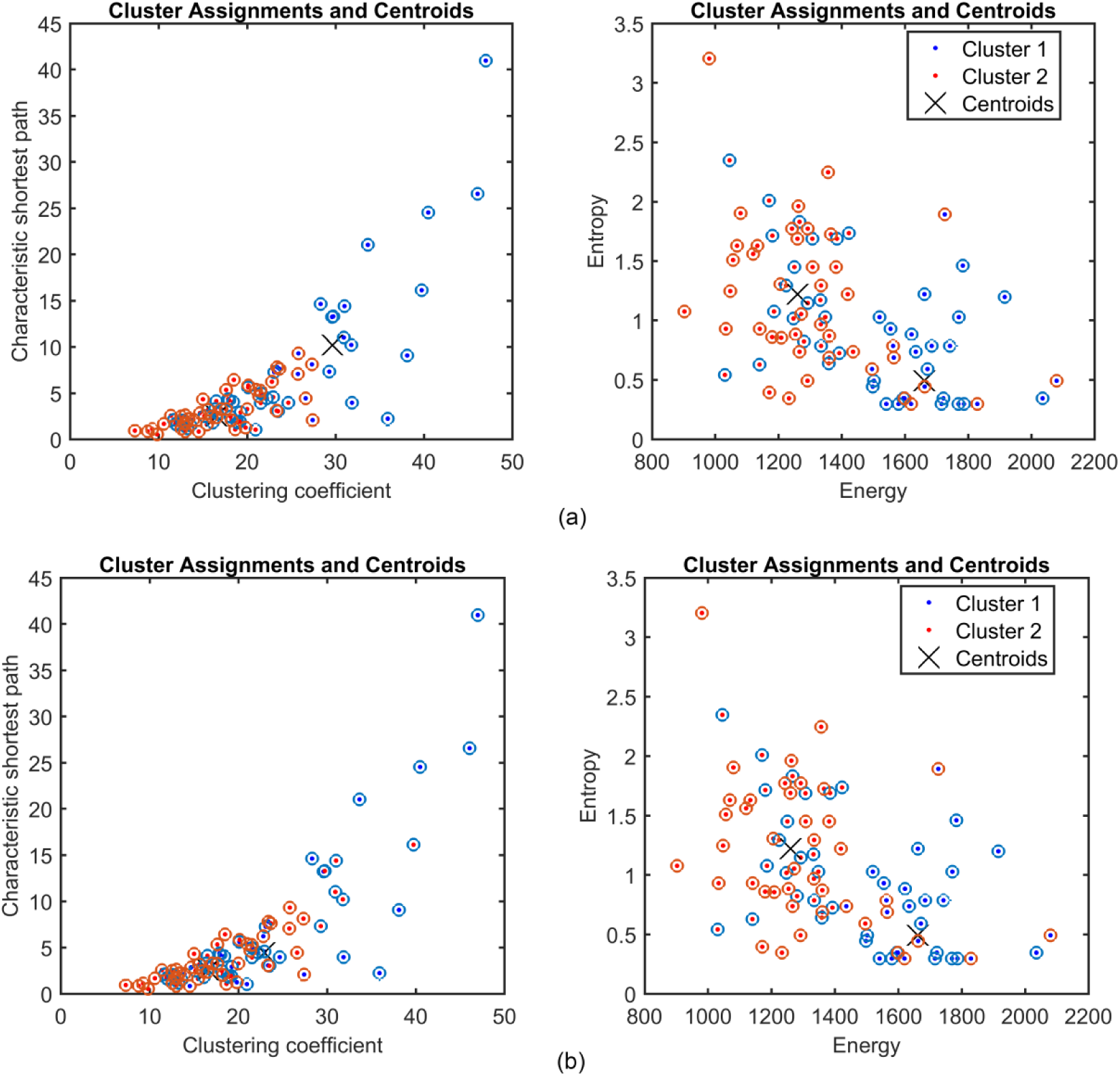
The scatter plot of clustering results by K-mean approach. Circles show individual values of graphs in the EC (blue circles) and the EO (red circles) condition. Dots show clusters that are assigned to each circle (individual data). If the color is matched between circle and dot (red-red or blue-blue), the clustering assignment is correct. **a)** Results of two-dimensional clustering based on the weighted graph measures (left side) and dynamical measures (right side) are shown in the scatter plot. b) Results of five-dimensional clustering based on the both topological weighted measures (clustering coefficient and characteristic shortest path) and dynamical measures (energy, entropy, largest eigenvalue). Both charts show the same clustering results but in different views (e.g. weighted dimensions (left side) and dynamical dimensions (right side)).

## 3. Discussion and conclusion

In this study, our aim was to investigate the overall link between different approaches to functional network connectivity analysis, using methods that have been previously investigated on their own or almost not at all. To this aim, we investigated topological and dynamical features of neural networks for two eye conditions as well as two random generated networks. Also, we performed Shannon entropy analysis, which we propose to be a plausible measure for evaluating the brain complexity. This would allow us to determine if there are any links between topology, synchrony and complexity of brain networks versus random networks. Our results indicate that: 1) the topological measures that we tested are effective in determining a state-dependent difference in brain connectivity and 2) brain synchrony can be best relayed by energy and largest eigenvalue. Lastly, there was no link between topology, synchrony and complexity of random networks however in the case of brain networks, several interactions can be found between these properties.

### 3-1 Differences between rsEC and rsEO conditions: Topological and spectral measures

#### 3-1-1 Topological measures

Clustering coefficient is closely associated with brain segregation, while brain integration is inversely related to characteristic shortest path ^7,10^. Based on our findings, the alpha network, that we explored here, exhibited higher segregation and lower integration in the rsEC condition compared with the rsEO condition. This result confirms the recent results of an EEG study that showed increased path length and clustering coefficient during rsEC condition in relation to the rsEO condition^48^. Therefore, it can be proposed that the alpha network tends to engage in more local processing when the eyes are closed versus open. Previous findings also indicate that a particular thalamocortical circuit that is specific to visual processing is involved in alpha band oscillation generation^49^. In particular, this oscillation band can be attributed to more posterior cortical regions^50^. Thus, it is no surprise that more segregation occurs in the rsEC condition than in the rsEO condition. However, seeing less integration in the EC condition is an emergent finding derived by the graph theory approaches described here.

However, topological measures associated with MST parameters, exhibited non-significant differences for the two (rsEO and rsEC) conditions. This result could be related to a lack of weak weights and loops as they are removed in the MST approach. In some literatures, the weak connections are considered in relation to noise ^51^. However, based on the nature of our connectivity measure (i.e. coherence) global noise (that can affect all the electrodes) can induce high value of coherence between electrodes. But, local noise (that can affect a few numbers of electrodes) may make weak connections between noisy electrodes and other electrodes. In this study, we performed a high-level control on artifact rejection and selected signals that were artifact free. Moreover, two conditions (rsEC and rsEO) were recorded at successive sessions, then possible local noise should affect both conditions. Then weak weights (that possibly cause significant differences between two conditions) cannot be considered in relation to noisy signals, and thus, may play a significant role in distinguishing the rsEC and rsEO conditions (at least when the connectivity measure is coherence).

#### 3-1-2 Spectral measures

Using spectral graph theory, we were able to distinguish between the two eye conditions in the brain network. Specifically, our results indicate that the rsEC condition exhibits higher energy than the rsEO condition. Previous findings have suggested that alpha activity is strongly synchronized in several electrodes especially posterior electrodes^50^. In literature, it has been shown that higher energy is directly related to a high degree of synchronization in the graph^52^. If we extend this to our findings, it follows that the higher energy of the rsEC condition predicts higher synchronization in comparison with the rsEO condition. Therefore, we can introduce energy as a powerful measure to find the level of synchronicity in functional brain networks.

Consistent with spectral graph theory literature^27,30,32^, we found that another spectral graph theory measure, the largest eigenvalue, can also be used to explore the synchronicity in functional brain networks. This measure also shows great sensitivity to the difference in the network between the two conditions. We suggest that this dynamical measure can be used in a reliable manner to ascertain the presence of differences in dynamical synchronization and desynchronization in resting state brain networks. Another spectral graph theory measure that we considered in this study was the second smallest eigenvalue, which was not significantly different between two conditions. The second smallest eigenvalue of graph is related to the robustness and stability of the network dynamic system^53^. There is not enough literature in network neuroscience to find the dynamical robustness and stability in functional brain networks during cognitive states (eyes closed or eyes open). Only a recent study has shown that Alzheimer’s disease decreases robustness of dynamic in the functional brain network^54^. This result, in addition to our result, can suggest that stability and robustness in dynamic of healthy brain remains constant during different cognitive states (at least during different states of visual processing), while diseases can change that.

### 3-2 Shannon entropy

The last measure that we supposed to be associated with the dynamic of functional brain network was entropy, and it was also significantly different between the two conditions. Specifically, higher entropy was observed in the rsEO condition compared with the rsEC condition. To frame this in terms of what is already known about entropy and network analysis, Shannon entropy describes the level of complexity and unpredictable information in a system (on a more general level) (Shannon, 1948). When measuring entropy in a network, higher entropy is attributed to a random and unpredictable network, whereas a network that has repeating or similar units is associated with the lowest entropy (i.e., an entropy of 0). In terms of our results, the network units (or the coherence between the electrodes in this case) in the rsEC condition are more predictable and simpler than for the rsEO condition. This suggests that there may be a decreased global synchronization of alpha band coherence in the rsEO condition compared with the rsEC condition. More synchronized systems (such as that associated with brain activity in the rsEC condition) exhibit similar correlations between coherence values. Shannon entropy may, thus, be another excellent tool to evaluate the synchronization (as well as complexity) of dynamical functional brain networks. Therefore, Shannon entropy (as well as energy and largest eigenvalue) can be utilized in the assessment or discrimination of different cognitive states.

### 3-3 Difference between brain-generated and random networks

Based on the results derived by calculation of L, C and entropy over brain generated networks vs. random networks, we did not see a specific pattern to distinguish these two types of networks. While the L, C and entropy in random exponential networks are in the order of brain-generated networks, the normal random network exhibits higher values for both L and C and lower value for entropy. It suggests that topological measures of a weighed graph and also entropy are more sensitive to distribution of weights and less sensitive to randomness of weight values. This is consistent with previous findings that suggested these measures are strongly related to distribution of weights and possible differences between networks may be affected by this bias^55^. Despite the measures of weighted graph, results indicate that MST measures are not dependent on distribution of weights but they are sensitive to randomness of weights. This result confirms a previous study that suggested MST as a weight-conserving approach ^55^. Same as MST measures, the spectral measures can clarify differences between brain-generated networks and random graphs. Both — the energy and the largest eigenvalue showed lower values for random networks. This suggests that EEG networks (brain-generated networks) are more synchronized than random networks with same mean and standard deviation of weights. Overall, these results suggest that topological properties of weighted networks are invariant to randomness of weights, but synchrony of networks differs from weight randomness.

### 3-4 Linking Entropy, Topological and Spectral Measures

One of our goals for this study was to ask whether topological and spectral measures are linked or not? To this aim, we performed the linear correlation analysis between measures that are derived from two kinds of networks (i. e. functional brain networks and random networks). In random networks (with normal and exponential distribution of weights), we found that measures are clustered independently from one another, suggesting that entropy, spectral, weighted and MST measures are in fact not linked.

However, there are various correlations between different types of measures in brain-generated networks. In both conditions (rsEC and rsEO), spectral measures (except second smallest eigenvalue), Shannon entropy and weighted graph measures are significantly correlated. There was no significant correlation (in rsEO condition) between second smallest eigenvalue (related to stability of dynamical networks) and other measures — especially energy and largest eigenvalue (associated with synchrony at dynamical networks). This result indicates that in low-level synchronized EEG network (rsEO), stability of network is independent from synchrony and topology of network. But in the high-synchronized EEG network (rsEC), stability is partly associated with synchrony (since it is significantly correlated with energy but not with largest eigenvalue). The correlation was positive, and it suggests high-level synchrony in EEG network can generate stability and robustness and vice versa.

Entropy is negatively correlated with spectral and weighted graph measures. The negative correlation between entropy and spectral measures suggests that brain networks with high synchrony have less complexity. The correlations between Shannon entropy and weighted graph measures can be considered in relation to segregation and integration in the EEG networks. Negative correlation between entropy and characteristic shortest path suggests the EEG network complexity is associated with integration within graph. On the other hand, negative entropy between entropy and clustering coefficient reveals complexity is inversely related to segregation in EEG networks. Mathematically, a regular graph (with same connectivity weights between nodes) presents minimum Shannon entropy, while, it exhibits maximum values for C and L^1^ and then it is not surprising to see the negative correlation between these measures. However, as demonstrated from our results, the negative association between Shannon entropy and C/L has not been observed in randomly generated graphs. Thus, our results suggest that this correlation is specifically distinguished in brain-generated networks (as a complex network).

Our results also revealed a positive correlation between the spectral and the weighted graph measures in brain-generated graphs (but not in random networks). These correlations suggest that, in the EEG networks, higher synchrony is associated with higher segregation (higher values for C) and lower integration (higher values for L). Therefore, one can conclude that local processing in EEG networks causes higher global synchrony, while global integration reduces level of synchrony in EEG networks. Mathematically, it can be justified by the argument that synchrony (energy and largest eigenvalue) is directly related to magnitude of weights but integration is reduced by higher values of weights (because L is directly associated with value of weights). This type of correlation has not been distinguished for random graph, as well as correlation between entropy and weighted graph measures. Then, it can be supposed that this type of correlation between spectral and topological graph measures is observed for specific types of complex networks, such as, EEG networks.

The last correlation measure for (brain-generated graphs) that we observed was between the weighted graph measures and the MST measures. This result was consistent with previous study that suggests same correlation^55^. As recommended by this study^55^ the mentioned correlations suggest both MST and weighted graph measures show same properties in topological brain networks, such as; segregation and integration.

In addition to the analysis of the individual parameters that are correlated within these sub-clusters, we also analyzed if we could compare the two eye condition states. We approached this problem using a k-means approach and we were able to determine that, when we utilize both topological and dynamical measures, we could more accurately differentiate between the networks associated with each of these cognitive states. A similar approach has been used once before, but only with topological measures (Ghaderi, Moradkhani, Haghighatfard, Akrami, Khayyer, & Balcı, 2018). Here, we showed that the extension of this method (also including dynamical measures) can prove to be a plausible approach in distinguishing different cognitive state-associated networks (Fig. 6).

### 3-5 Finding the best measure for the EEG network

While revealing brain networks related to the variable that is being tested (in our case, the cognitive state related to whether the eyes are open or closed), it is important to consider which measures are capable of distinguishing between these different states. However, it is more intricate than that. A good measure has to meet three important criteria: 1) it must be able to distinguish between variable-related and randomly generated data; 2) it must be independent of the distribution of weights (see ^55^ for discussion); and 3) it must be able to differentiate between the levels of the variable being tested (i.e., between the rsEO and rsEC condition networks here).

From our results, we were able to determine the measures that have met these criteria. Importantly, only one measure met all three criteria: largest eigenvalue. Previously, weighted graph measures were used to distinguish variable-related states ^7,10^. Later it has also been proposed that MST measures could be used to accomplish this goal^55,56^. Here, we showed that weighted graph measures met the third criterion only, whereas most MST measures met the first two criteria. Taken together, we suggest that, when it is important to distinguish between dynamical networks (such as EEG network) that must satisfy all three of these criteria, largest eigenvalue could be used; otherwise, weighted graph and MST measures should be applied based on the nature of the variable being tested.

We also proposed three measures (energy, largest eigenvalue, and entropy) as possible measures that could distinguish between rsEO and rsEC condition-related networks. However, from these three measures, energy met two of the three criteria, whereas entropy only met one criterion. Energy was able to distinguish between variable-related and random data, differentiate between rsEO and rsEC condition networks, but, it was sensitive to the weight distribution. On the other hand, entropy was only sensitive to rsEO versus rsEO condition networks, which is a similarity between entropy and other weighted graph measures.

Overall, these considerations suggest that it is important to consider the nature of the independent variable being tested, as well as which criteria need to be met. Certain analyses may require that only some or particular criteria need to be met, and thus, the appropriate measure(s) should be chosen, be it weighted graph, MST, or those proposed here, based on the requirements of the network.

### 3-6 Conclusion

In this study, we investigated and introduced new measures to distinguish different types of networks (two brain-generated and two random generated networks). Not only did we investigate the ability of topological measures to make this distinction, but we also introduced dynamical measures in this endeavor. We showed here that topological and dynamical measures exhibited different properties of networks. These two types of measures were completely independent in the case of random-generated networks, however, several significant correlations between topology and dynamic of EEG network (brain generated networks) were found. To distinguish between two types of brain networks, topological measures show that alpha bands are more segregated in the rsEC condition and are more integrated in the rsEO condition. On the other hand, dynamical measures suggest that the alpha network is more synchronized in the rsEC condition. In terms of complexity, Shannon entropy showed that the complexity of the rsEO state is higher than for the rsEC state. Shannon entropy may, thus, be used to investigate the level of complexity in a functional brain network. Having both topological and dynamical measures made it more helpful to separate the two cognitive state-related networks that we investigated (using a k-means approach). Thus, the methodology used in this study helps to provide information about the synchronicity and complexity of brain networks. However, to achieve these analyses, it is especially important to consider which measures are best suited to distinguish between such state-related networks (i.e., sensitivity to random networks, distribution, and ability to separate between cognitive-related states). Of all the measures that we investigated, we found that the largest eigenvalue is a great candidate for this goal, as it met all three criteria. Overall, further investigations of all these measures in functional network analysis are required, as the findings may have major implications for many cognitive and clinical studies.

## 4 Method

### 4-1 Participants, ethical considerations and EEG acquisition

We reanalyzed a previous open access dataset (https://figshare.com/articles/Over_zip/5970886) from the same participants in the rsEC and rsEO conditions. The ethical processes and EEG acquisition/data were similar to our previous published paper^14^. The details about ethical considerations, participants and EEG recordings are in the previously published work^14^.

### 4-2 Pre-analysis of EEG data

EEG activity was recorded for five minutes for each condition (rsEO and rsEC). Artifacts were rejected using a z-score-based algorithm (NeuroGuide software; www.appliedneuroscience.com). Signal selection was then performed manually by an expert. Since the signal-length plays a significant role in calculating the value of connectivity measures^57^ we used a similar signal-length for all participants and conditions. In each condition, 20 to 25 artifact-free signal segments with a length of three seconds were selected for fast Fourier transform (FFT) analysis. This analysis was performed separately for each segment with a 25 % sliding window ^58^ for FFT analysis. As alpha rhythm is more affected by eye conditions^48,59^, we applied a bandwidth filter (8 to 12 Hz, i.e., alpha rhythm) on the original signal and then alpha band was used for further analysis.

### 4-3 EEG connectivity and adjacency matrix

Although high-density EEG studies with more than 64 recording channels can display a highly spatial resolution of brain function and the number of electrodes or nodes can improve the GTA analysis, several recent studies with limited recording channels indicate that reliable results can also be obtained using standard 10/20 system with 19 EEG channels^12,13, 60–63^. In order to compute a functional brain network, we used coherence as a simple and well-studied EEG connectivity measure ^4,12,14, 64–67^. Although several criticisms (e.g., effect of volume conduction) have been raised against coherence^68^, still it is a useful and meaningful measure when coupling and synchronizing between neural units are important^64,69–71^. Furthermore, when topological measures were investigated, a recent study suggested that *coherence* might be a better measure than *phase order parameter* and *synchronization likelihood* to expose more significant differences between groups or conditions^65^. On the other hand, the effect of volume conduction is more important when EEG electrodes are located very close together (high-density EEG). But in 19-channel system (as we used in this study), the reliability of 1×N connectivity derived by coherence is greater than phase synchrony estimates^72^.

We used the function ‘*mscohere’* in MATLAB to find magnitude-squared coherence in alpha band (8-12 Hz). Mathematically this function calculates ^73^:

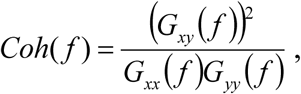

where *G_xy_* (*f*) is a cross-spectrum between signals x and y and *G_xx_*(*f*), *G_yy_*(*f*) are auto-spectra for x and y respectively. In this equation, coherence is related to the phase difference between signals *x* and *y*. When the phase changes randomly between signals, then coherence is close to ‘zero’, whereas coherence is equal to ‘one’ when the phase is constant between signals^74^. In this study, coherence was calculated between 171 pairs 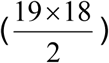 of channels which was then used to generate the adjacency matrix. In the adjacency matrix, each row/column represents a channel and their intersection in the matrix corresponds to the coherence between channels. This weighted adjacency matrix was subsequently used for further analysis (as described in the following sections).

### 4-4 Topological indices

#### 4-4-1 Clustering coefficient and characteristic shortest path of weighed graphs

We used two well-studied measures (i. e., clustering coefficient (*C*) and characteristic shortest path (*L*)) to investigate the topology of weighted graphs ^7,8,10^. The clustering coefficient helps to identify features related to the level of segregation in the network^10^. The *C* of the node *‘i’* in an undirected weighted network, such as that in our study is defined as^75^:

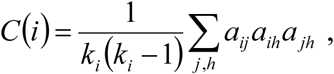

where *k_i_* is the degree of the node *‘i’* and *a_ij_* defines the connectivity weight between the nodes *‘i’* and *‘j’*.

Another index of a weighted graph that can be determined is the characteristic shortest path^10^, which uncovers information about the integration in a complex brain network. In an undirected weighted graph, the shortest path between two nodes *‘i’* and *‘j’* is measured by the minimum summation of weights between those two nodes, where the characteristic shortest path is the average of all the shortest paths between all possible pairs of nodes^10^:

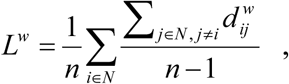

Where 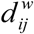 is the distance between two nodes *‘i’* and *‘j’* and *‘n’* is number of the paths in the graph.

To calculate these measures, we used the matrices obtained in the previous section that contain coherence values for each combination of the 19 pairs of electrodes. These matrices were calculated for both the rsEO and rsEC conditions (i.e., two matrices (rsEO, rsEC) for each participant) as well as random matrices with normal and exponential distributions. We have used the *brain connectivity toolbox*^10^ to calculate the clustering coefficient and characteristic shortest path of weighed graphs.

#### 4-4-2 Minimum Spanning Tree (MST) analysis

Typically, MST finds a backbone (tree) for a weighted graph^56^. In the weighted networks, nodes can be connected through multiple paths to each other and several loops may be generated via a graph. These loops and multiple paths lead to graph measure dependency on factors such as the average of connection power and the distribution of weights^51,55^. To investigate changes of the MST measures according to weight distribution and randomness of weights, and to evaluate the correlations between MST measures and SGT measures we performed MST analyses for two experimental conditions (rsEO and rsEC) vs. two randomly generated dataset with different weight distributions.

To compare these matrices, four MST indices were calculated: 1) betweenness centrality, 2) leaf fraction, 3) diameter, and 4) eccentricity. Betweenness centrality of a node is defined as the number of paths (specifically shortest paths) that emerge from a specific node. In the MST approach, all the paths between nodes are the shortest path (because no loops are possible), which allows us to define betweenness centrality simply as a function of all the paths that cross a particular node. This allows us to report a betweenness centrality measure for all the nodes in a network; however, we can also average this measure across all nodes for a particular condition and report a single value only. The more integrated the graph, nodes will exhibit higher betweenness centrality^10^.

In the MST approach, the second index is the leaf fraction which is associated with the number of nodes at the end of the chain. Here, a chain is known as a particular set of connections of nodes in a network from the beginning to the end. Within the chain, a leaf is defined as a node that has a degree equal to a value of ‘1’. Mathematically, the leaf fraction is equal to the number of nodes with a degree of value equal to ‘1’ (*N_(k=1)_*) divided by *N-1*, where *N* represents the total number of nodes in a tree^51^. When the tree has a central node connected to all other nodes, the value of the leaf fraction is maximal; on the other hand, the leaf fraction is at a minimum for a tree with a line shape where one node is connected uniquely to another node in the network ^13^. In terms of integration of a network, trees with a network in the shape of a line that have low leaf fraction value are deemed to be less integrated than graphs with a flower shape and high leaf fraction value ^56^.

The maximum path length in a tree defines the third MST index which is the diameter. This is same as the shortest path measure in a weighted graph. Diameter is inversely proportional to the level of integration in a network, such that a longer path is associated with less integration and a shorter path is reflective of greater integration ^51^.

The final MST index is eccentricity, which is defined as the longest distance of a node to any other node in either direction of a chain. Each node can be assigned an eccentricity, but we can also calculate an average eccentricity for the entire tree. In a tree, eccentricity is negatively related to integration, whereby graphs with a higher average eccentricity have highly isolated nodes and show less integration, and vice versa. First, we used the biograph toolbox in MATLAB (ver. 2016a) to compute the MST tree of weighted graphs, the results of which we subsequently fed into the brain connectivity toolbox ^10^ to obtain the final four MST indices.

## 5 Dynamical indices

### 5-1 Spectral graph theory (eigenvalues and energy)

Mathematically, each oscillatory system with different units of oscillators can be presented by a symmetric connectivity matrix (A) of oscillations between units (e.g., an EEG channel) and each symmetric connectivity matrix can be decomposed into a set of eigenvalues and eigenvectors that follow the bellow equation:

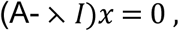

where *I* is a single unit matrix, *x* is eigenvector of A and ⋋ is the eigenvalue matrix, if and only if Det |A- ⋋ *I* | = 0.

In graph theory, eigenvalue spectrums associated with adjacency or Laplacian (i.e., adjacency matrix minus unit matrix of degrees associated with the nodes in a network) matrices are commonly considered as measures related to dynamics and synchronization of a network^76^. In SGT in particular, the largest eigenvalue and energy are considered as two important measures related to synchrony^27,29,77^ and the stability of the network^78^. The energy of a graph is defined as the sum of the absolute values of eigenvalues^79^. This energy is directly related to the number of nodes and value of edges ^79^. Therefore, it is expected that, in a dynamical system with constant nodes (e.g., 19 in this study for each of the 19 electrodes), higher energy would be achieved when the synchronization or coupling is increased in the graph^79^. Energy is defined by:

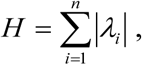

where *γ_i_* is the eigenvalue spectrum and *n* is the number of nodes. Since the adjacency matrices of brain networks are typically symmetric and positive, the SGT approach can be easily applied to them. As a result, the eigenvalues of these networks are always computable and non-negative. We compared largest eigenvalue and energy between all conditions (e.g. rsEC, rsEO, normal distribution, exponential distribution). We took our adjacency matrices for these conditions and then applied the *eig* function in MATLAB (ver. 2016a) to these matrices in order to calculate eigenvalues.

### 5-2 The Shannon entropy of brain network

Entropy measures have been employed to investigate functional brain connectivity. Transfer entropy^80^ and mutual information^81^ are appropriate measures to evaluate the level of disorder between two signals. However, although these measures can demonstrate the complexity of synchronization/desynchronization between two neural regions or electrodes, they cannot evaluate the complexity of global brain networks. To study the global complexity of a network, the entropy of graph degrees^3^ and graph spectrum^82^ has been proposed as two valid measures. It is important to note that the network entropies that focus on the complexity of the overall network require a relatively large number of nodes (e.g. N≈100). However, the limited number of electrodes in many EEG setups (same as our setup) precludes the use of such measures. Thus, in order to investigate the complexity of a brain network, given this limitation, we used Shannon or information entropy ^83^. Mathematically, Shannon entropy is defined by:

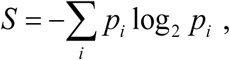

where *p* is the probability of *i* (the possible values within an array in the adjacency matrix). According to this equation, a system with maximum Shannon entropy (with fully random behavior) has non-repetitive values in the array, while Shannon entropy is equal to zero in a deterministic system (e.g., matrix of ones). The number of possible values of array cells (i.e., histogram count) is directly related to the Shannon entropy^84^.

In this study, we considered Shannon entropy for a 19×19 adjacency matrix. We introduced the application of Shannon entropy of an adjacency matrix to provide *information entropy of brain network*. This measure is suggested for the use of evaluating the complexity of global brain connectivity. We calculated this measure for a weighted adjacency matrix derived by short term signals (i.e., duration on the order of seconds). We used the *Entropy* function in the image processing toolbox of MATLAB (ver. 2016a) that works based on the *Imhist* function.

## 6 Statistical analysis

### 6-1 Probability distribution of an adjacency matrix

The probability distributions of the adjacency matrix were considered in the rsEO and rsEC conditions. To this end, we constructed 45 adjacency matrices with a normal distribution and 45 adjacency matrices with a random exponential distribution. We used the mean and standard deviation of brain-generated adjacency matrices to construct the random values. The probability distribution was presented through histograms. We used 200 bins for each histogram between 0 and 100 (the values of coherence are between 0 and 1 and we multiplied all the coherence values by 100 for the ease of visualization and analysis). This number of bins results in an appropriate representation of the data.

Then, the r-squared errors between distributions were calculated which were ultimately used to determine the distances between probability distributions. We used this probability distribution analysis to find the effects of probability distribution on dynamical or statistical graph measures.

### 6-2 Permutation test

A nonparametric permutation test^85^ with 10,000 random shuffles was used to compare graph measures between two conditions. To avoid *type 1* error, a false discovery rate (FDR) analysis ^86^ was performed and then corrected p-values (q-values) were obtained. We performed FDR separately for weighted graph measures (i.e., clustering coefficient, characteristic shortest path), MST measures (i.e., betweenness centrality, leaf fraction, eccentricity, and diameter) and dynamical measures (i.e., largest eigenvalue, energy and entropy). The permutation test was carried out using a previously created MATLAB function^13,14^. After that, an individual comparison was performed to identify significant measures between the two conditions. We also used the FDR analysis to correct multiple comparisons between conditions (e. g. rsEC, rsEO, random exponential, random normal).

### 6-3 Correlation analysis and K-means

Correlation analysis was performed between all measures using the MATLAB function *corr* and then the p-values were obtained. To avoid the effect of multiple comparisons, Bonferroni correction was performed. After that, a K-means clustering approach was used to separate the data into two clusters. Topological measures (i.e., clustering coefficient and shortest path of a weighted graph), MST measures, and dynamical measures (i.e., energy, max eigenvalue and entropy) were used separately for clustering, as well as a combination of these measures. All these factors and combinations, thereof, were considered in order to determine the best measures at clustering the data. The accuracy of clustering was then compared in order to determine the most appropriate measure to cluster the data in a linear manner. All statistical analyses were performed in MATLAB (ver. 2016a), and the *ttest1* function (for within subject t-test analysis) was used specifically to compare the means of the conditions.

